# Text-mining clinically relevant cancer biomarkers for curation into the CIViC database

**DOI:** 10.1101/500686

**Authors:** Jake Lever, Martin R. Jones, Arpad M. Danos, Kilannin Krysiak, Melika Bonakdar, Jasleen Grewal, Luka Culibrk, Obi L. Griffith, Malachi Griffith, Steven J.M. Jones

**Affiliations:** Canada’s Michael Smith Genome Sciences Centre, Vancouver, BC, Canada; University of British Columbia, Vancouver, BC, Canada; McDonnell Genome Institute, Washington University School of Medicine, St. Louis, MO, USA; Siteman Cancer Center, Washington University School of Medicine, St. Louis, MO, USA; Division of Oncology, Department of Medicine, Washington University School of Medicine, St. Louis, MO, USA; Department of Genetics, Washington University School of Medicine, St. Louis, MO, USA; Simon Fraser University, Burnaby, BC, Canada

## Abstract

Precision oncology involves analysis of individual cancer samples to understand the genes and pathways involved in the development and progression of a cancer. To improve patient care, knowledge of diagnostic, prognostic, predisposing and drug response markers is essential. Several knowledgebases have been created by different groups to collate evidence for these associations. These include the open-access Clinical Interpretation of Variants in Cancer (CIViC) knowledgebase. These databases rely on time-consuming manual curation from skilled experts who read and interpret the relevant biomedical literature. To aid in this curation and provide the greatest coverage for these databases, particularly CIViC, we propose the use of text mining approaches to extract these clinically relevant biomarkers from all available published literature. To this end, a group of cancer genomics experts annotated biomarkers and their clinical associations discussed in 800 sentences and achieved good inter-annotator agreement. We then used a supervised learning approach to construct the CIViCmine knowledgebase (http://bionlp.bcgsc.ca/civicmine/) extracting 128,857 relevant sentences from PubMed abstracts and Pubmed Central Open Access full text papers. CIViCmine contains over 90,992 biomarkers associated with 7,866 genes, 402 drugs and 557 cancer types, representing 29,153 abstracts and 40,551 full-text publications. Through integration with CIVIC, we provide a prioritised list of curatable biomarkers as well as a resource that is valuable to other knowledgebases and precision cancer analysts in general.

## Introduction

The ability to stratify patients into groups that are clinically related is an important step towards a personalized approach to cancer. Over time, a growing number of biomarkers have been developed in order to select patients who are more likely to respond to certain treatments. These biomarkers have also been valuable for prognostic purposes and for understanding the underlying biology of the disease by defining different molecular subtypes of cancers that should be treated in different ways (e.g. *ERBB2*/*ESR1*/*PGR* testing in breast cancer [1]). Immunohistochemistry techniques are a primary approach for testing samples for diagnostic markers (e.g. CD15 and CD30 for Hodgkin’s disease [2]). Recently, the lower cost and increasing speed of genome sequencing has also allowed the DNA and RNA of individual patient samples to be characterized for clinical applications [3]. Throughout the world, this technology is beginning to inform clinician decisions on which treatments to use [4]. Such efforts are dependent on comprehensive and current understanding of the clinical relevance of variants. For example, the Personalized Oncogenomics project at BC Cancer identifies somatic events in the genome such as point mutations, copy number variations and large structural changes and, in conjunction with gene expression data, generates a clinical report to provide an ‘omic picture of a patient’s tumor [5].

The high genomic variability observed in cancers means that each patient sample includes a large number of new mutations, many of which may have never been documented before [6]. The phenotypic impact of most of these mutations is difficult to discern. This problem is exacerbated by the driver/passenger mutation paradigm where only a fraction of mutations are essential to the cancer (drivers) while many others have occurred through mutational processes that are irrelevant to the progression of the disease (passengers). An analyst trying to understand a patient sample typically performs a literature review for each gene and specific variant which is needed to understand its relevance in a cancer type, characterize the driver/passenger role of its observed mutations, and gauge the relevance for clinical decision making.

A number of groups have built in-house knowledgebases, which are developed as analysts examine increasing numbers of cancer patient samples. This tedious and largely redundant effort represents a substantial interpretation bottleneck impeding the progress of precision medicine [7]. To encourage a collaborative effort, the CIViC knowledgebase (https://civicdb.org) was launched to provide a wiki-like, editable online resource where community contributed edits and additions are moderated by experts in order to maintain high quality variant curation[8]. The resource provides information about clinically-relevant variants in cancer described in the peer-reviewed literature. Variants include protein-coding point mutations, copy number variations, epigenetic marks, gene fusions, aberrant expression levels and other ‘omic events. It supports four types of evidence associating biomarkers with different classes of clinical relevance (also known as evidence types).

Diagnostic evidence items describe variants that can help a clinician diagnose or exclude a cancer. For instance, the *JAK2* V617F mutation is a major diagnostic criterion for myeloproliferative neoplasms to identify polycythemia vera, essential thrombocythemia and primary myelofibrosis [9]. Predictive evidence items describe variants that help predict drug sensitivity or response and are valuable in deciding further treatments. Predictive evidence items often explain mechanisms of resistance in patients who progressed on a drug treatment. For example, the *ABL1* T315I missense mutation in the *BCR-ABL* fusion, predicts poor response to imatinib, a tyrosine kinase inhibitor that would otherwise effectively target *BCR-ABL*, in patients with chronic myeloid leukemia [10]. Predisposing evidence items describe germline variants that increase the likelihood of developing a particular cancer, such as *BRCA1* mutations for breast/ovarian cancer [11] or *RB1* mutations for retinoblastoma [12]. Lastly, prognostic evidence items describe variants that predict survival outcome. As an example, colorectal cancers that harbor a *KRAS* mutation are predicted to have worse survival [13].

CIViC presents this information in a human-readable text format consisting of an ‘evidence statement’ such as the sentence describing the ABL1 T315I mutation above together with data in a structured, programmatically accessible format. A CIViC ‘evidence item’ includes this statement, ontology-associated disease name [14], evidence type as defined above, drug (if applicable), PubMed ID and other structured fields. Evidence items are manually curated and associated in the database with a specific gene (defined by Entrez Gene) and variant (defined by the curator).

Several groups have created knowledgebases to aid clinical interpretation of cancer genomes, many of whom have joined the Variant Interpretation for Cancer Consortium (VICC, http://cancervariants.org/). VICC is an initiative that aims to coordinate variant interpretation efforts and, to this end, has created a federated search mechanism to allow easier analysis across multiple knowledgebases [15]. The CIViC project is co-leading this effort along with OncoKB [16], the Cancer Genome Interpreter [17], Precision Medicine Knowledge base [18], Molecular Match, JAX-Clinical Knowledge base [19] and others.

Most of these projects focus on clinically-relevant genomic events, particularly point mutations, and provide associated clinical information tiered by different levels of evidence. Only CIViC includes RNA expression-based biomarkers. These may be of particular value for childhood cancers which are known to be ‘genomically quiet’, having accrued very few somatic mutations. Consequently, their clinical interpretation may rely more heavily on transcriptomic data [20]. Epigenomic biomarkers will also become more relevant as several cancer types are increasingly understood to be driven by epigenetic misregulation early in their development [21]. For example, methylation of the MGMT promoter is a well known biomarker in brain tumors for sensitivity to the standard treatment, temozolomide [22].

The literature on clinically relevant cancer mutations is growing at an extraordinary rate. For instance, only 5 publications in PubMed mentioned BRAF V600E in the title or abstract in 2004 compared to 454 papers in 2017. In order to maintain a high quality and up-to-date knowledge-base, a curation pipeline must be established. This typically involves a queue for papers, triaging those that should be curated and then assignment to a highly experienced curator. This prioritisation step is important given the limited time of curators and the potentially vast number of papers to be reviewed. Prioritisation must identify papers that contain knowledge that is of current relevance to users of the knowledgebase. For instance, selecting papers for drugs that are no longer clinically approved would not be valuable to the knowledgebase.

Text mining methods have become a common approach to help prioritise literature curation. These methods fall broadly into two main categories, information retrieval (IR) and information extraction (IE). IR methods focus on paper-level information and can take multiple forms. Complex search queries for specific terms or paper metadata (helped by the MeSH term annotations of papers in biomedicine) are common tools for curators. More advanced document clustering and topic modelling systems can use semi-supervised methods to predict whether a paper would be relevant for curation. Examples of this approach include the document clustering method used for the ORegAnno project [23].

IE methods extract structured knowledge directly from the papers. This can take the form of entity recognition, by explicitly tagging mentions of biomedical concepts such as genes, drugs and diseases. A further step can involve relation extraction to understand the relationship discussed between tagged biomedical entities. This structured information can then be used to identify papers relevant for the knowledgebase. IE methods are also used for automated knowledgebase population without a manual curation step. For example, the miRTex knowledgebase, which collates microRNAs and their targets, uses automated relation extraction methods to populate the knowledgebase [24]. Protein-protein interaction networks (such as STRING [25]) are often built using automatically generated knowledgebases.

The main objective of this project was to identify frequently discussed cancer biomarkers that fit the CIViC evidence model but are not yet included in the CIViC knowledgebase. We developed an IE-based method to extract key parts of the evidence item: cancer type, gene, drug (where applicable) and the specific evidence type from published literature. This allows us to count the number of mentions of specific evidence items in abstracts and full text articles and compare against the CIViC knowledgebase. This paper will present our methods to develop this resource, known as CIViCmine (http://bionlp.bcgsc.ca/civicmine/). The main contributions of this work are an approach for knowledgebase construction that could be applied to many areas of biology and medicine, a machine learning method for extracting complicated relationships between four entity types, and extraction of relationships across the largest possible publically-accessible set of abstracts and full text articles. This resource, containing 90,992 gene-cancer associations with clinical relevance, is valuable to all cancer knowledgebases to aid their curation and also as a tool for precision cancer analysts searching for evidence supporting biomarkers not yet included in any other resource.

## Methods

### Corpora

The full PubMed and PubMed Central Open Access subset corpora was downloaded from the NCBI FTP website using the PubRunner infrastructure [26]. These documents were converted to the BioC format for processing with the Kindred package [27]. HTML tags were stripped out and HTML special characters converted to Unicode. Metadata about the papers were retained including PubMed IDs, titles, journal information and publication date. Subsections of the paper were extracted using a customised set of acceptable section headers such as “Introduction”, “Methods”, “Results” and many synonyms of these (accessible through the GitHub repository). The corpora were downloaded in bulk in order to not overload the EUtils RESTFUL service that is offered by the NCBI. In order to avoid duplications of publications in PMCOA and PubMed, the PMIDs of all documents included in PMCOA were used to filter out abstracts from the PubMed corpus. The updated files from PubMed were also processed to identify the latest version of each abstract to process.

### Term Lists

Term lists were curated for genes, diseases and drugs based on several resources. The cancer list was curated from a section of the Disease Ontology [14]. All terms under the “cancer” (DOID:162) parent term were selected and filtered for nonspecific names of cancer (e.g. “neoplasm” or “carcinoma”). These cancer types were then matched with synonyms from the Unified Medical Language System (UMLS) Metathesaurus [28] (2017AB), either through existing external reference links in the Disease Ontology or through exact string-matching on the main entity names. The additional synonyms in the UMLS were then added through this link. The genes list was built from the Entrez gene list and complemented with UMLS terms. Terms that overlapped with common words found in scientific literature (e.g. ice) were removed.

The drug list was curated from the WikiData resource [29]. All Wikidata entities that are drug instances (Wikidata identifier: Q12140) were selected using a SPARQL query. The generic name, brand name and synonyms were extracted where possible. This list was complemented by a custom list of general drug categories (e.g. chemotherapy, tyrosine kinase inhibitors, etc) and a list of inhibitors built using the previously discussed gene list. This allowed for the extraction of terms such as “EGFR inhibitors”. This was done because analysts are often interested in and publications often discuss biomarkers associated with drug classes that target a specific gene, in addition to specific drugs.

All term lists were filtered with a stopwords list. This was based on the stopword list from the Natural Language Toolkit [30] and the most frequent 5,000 words found in the Corpus of Contempory American English [31] as well as a custom set of terms. It was then merged with common words that occur as gene names (such as ICE).

A custom variant list was built that captured the main types of point mutations (e.g. loss of function), copy number variation (e.g. deletion), epigenetic marks (e.g. promoter methylation) and expression changes (e.g. low expression). These variants were complemented by a synonym list.

The word lists and tools used to generate them are accessible through the BioWordlists project (https://github.com/jakelever/biowordlists) and data can be found in the Zenodo repository ((http://doi.org/10.5281/zenodo.1286661).

### Entity extraction

The BioC corpora files were processed by the Kindred package. This NLP package used Stanford CoreNLP [32] for processing in the original published version [27]. For this work, it was changed to Spacy [33] for the improved Python bindings in version 2 for this project. This provided easier integration and execution on a cluster without running a Java subprocess. Spacy was used for sentence splitting, tokenization and dependency parsing of the corpora files.

Exact string matching was then used against the tokenized sentences to extract mentions of cancer types, genes, drugs and variants. Longer terms were prioritised during extraction so that “non small cell lung cancer” would be extracted instead of just “lung cancer”. Variants were also extracted with a regular expression system for extracting protein coding point mutations (e.g. V600E).

Gene fusions (such as *BCR-ABL1*) were detected by identifying mentions of genes separated by a forward slash, hyphen or colon. If the two entities had no overlapping HUGO IDs, then it was flagged as a possible gene fusion and combined into a single entity. If there were overlapping IDs, it was deemed likely to be referring to the same gene. An example is *HER2/neu* which is frequently seen and refers to a single gene (*ERBB2*) and not a gene fusion.

Acronyms were also detected, where possible, by identifying terms in parentheses and checking the term before it, for instance “non-small cell lung carcinoma (NSCLC)”. This was done to remove entity mistakes where possible. The acronym detection method takes the short form (the term in brackets) and iterates backwards through the long form (the term before brackets) looking for potential matches for each letter. If the long form and short form has overlapping associated ontology IDs, they likely refer to the same thing and can be combined, as in the example above. If only one of the long form or short form has an associated ontology ID, they are combined and assigned the associated ontology ID. If both long form and short form have ontology IDs but there is no overlap, the short form is disregarded as the long form has more likelihood of getting the specific term correct.

Gene mentions that are likely associated with signalling pathways and not specific genes (e.g. “MTOR signalling”) are also removed using a simple pattern based on the words after the gene mention. One final post-processing step merges neighbouring terms with matching terms. So “HER2 neu” would be combined into one entity as the two terms (*HER2* and *neu*) refer to the same gene.

### Sentence selection

With all biomedical documents parsed and entities tagged, all sentences were selected that mention at least one gene, at least one cancer and at least one variant. A drug was not required as only one (predictive) of the four evidence types involves a drug entity. These sentences were enriched by filtering with certain keywords that are strongly associated with the different evidence items. The full list and groupings of keywords are shown in Table 1). This grouping is done to make sure that each evidence type is represented reasonably equally in the training data. The General category with the keyword “marker” is included to catch additional sentences that discuss markers, which may relate to any of the four evidence types. Several of the keywords are stems in order to capture different forms of the word, e.g. prognosis or prognostic. The acronym “DFS” which means “disease free survival” is also included as it was found in many sentences describing prognosis.

**Table 1.**
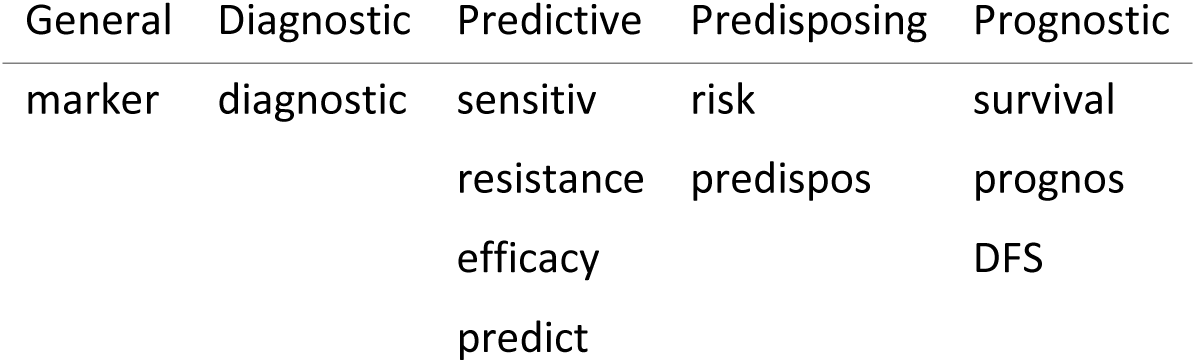
The five groups of search terms used to identify sentences that potentially discussed the four evidence types. Strings such as “sensitiv” are used to capture multiple words including “sensitive” and “sensitivity”.

| General | Diagnostic | Predictive | Predisposing | Prognostic |
| --- | --- | --- | --- | --- |
| marker | diagnostic | sensitiv | risk | survival |
|  |  | resistance | predispos | prognos |
|  |  | efficacy |  | DFS |
|  |  | predict |  |  |

### Annotation Platform

A web platform for simple relation annotation was built using Bootstrap (https://getbootstrap.com/). This allowed annotators to work using a variety of devices, including their smartphones. The annotation system could be loaded with a set of sentences with entity annotations stored in a separate file (also known as standoff annotations). When provided with a relation pattern, for example “Gene/Cancer”, the system would search the input sentences and find all pairs of the given entity types in the same sentence. It would make sure that the two entities are not the same term, as in some sentences a token (or set of tokens) could be annotated as both a gene name and a cancer type (e.g. “retinoblastoma”). For a sentence with 2 genes and 2 cancer types, it would find all four possible pairs of gene and cancer type.

Each sentence, with all the possible candidate relations matching the relation pattern, would be presented to the user, one at a time (Fig 1). The user can then select various toggle buttons for the type of relation that these entities are part of. They can also use these to flag entity extraction errors or mark contentious sentences for discussion with other annotators.

**Figure 1.**
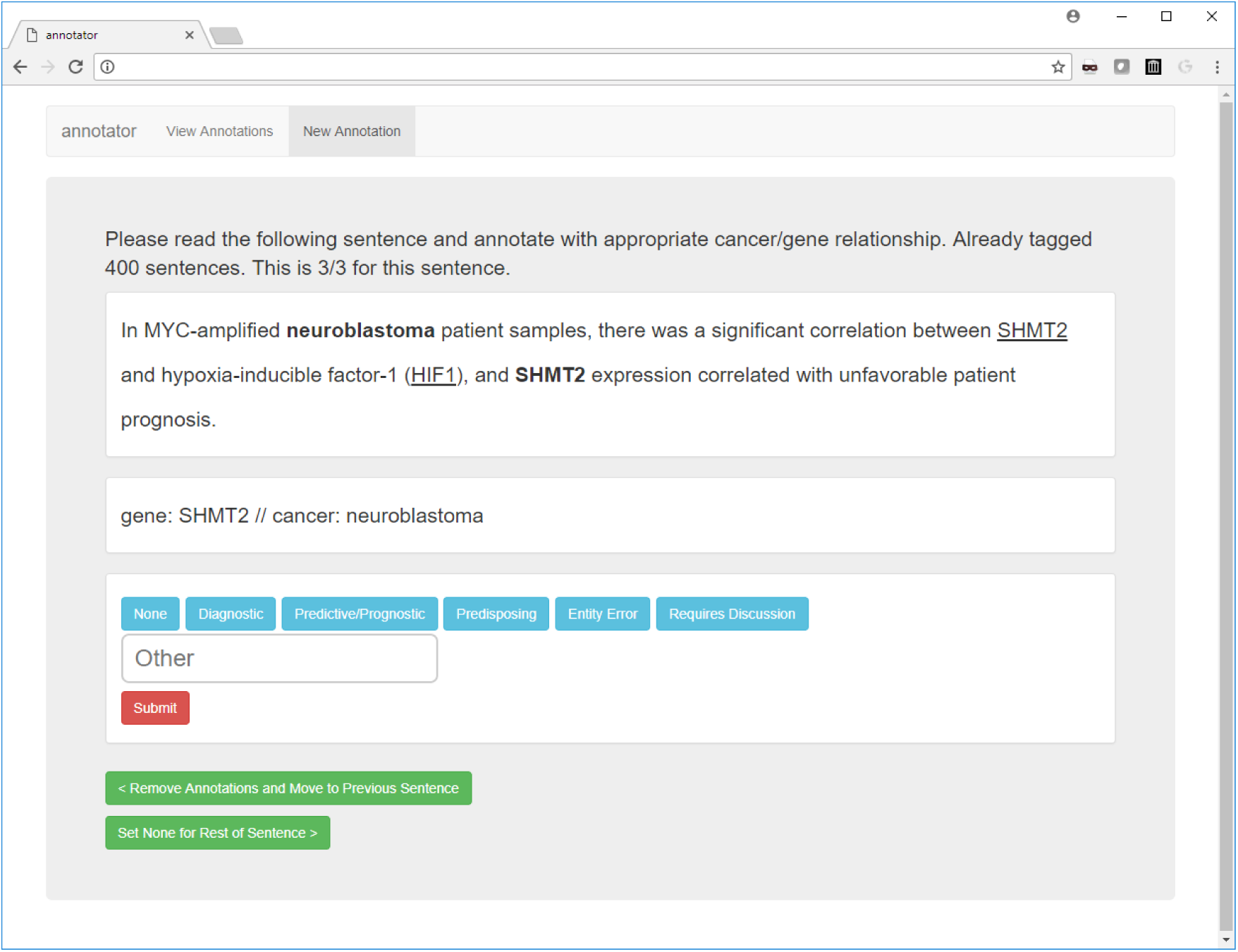
A screenshot of the annotation platform that allowed expert annotators to select the relation types for different candidate relations in all of the sentences. The example sentence shown describes a prognostic marker.

### Annotation

For the annotation step (outlined in Fig 2), the annotated data set (known as the gold set) was constructed using a consensus of multiple annotators. An equal number of sentences were selected from each of the groups outlined in Table 1. This guaranteed coverage of all four evidence types as otherwise the prognostic type dominated the other groups. If this step was not done, 100 randomly selected sentences would only contain 2 (on average) from the diagnostic group. However, this sampling provided poor coverage of sentences that describe specific point mutations. Many precision oncology projects only focus on point mutations and so a further requirement was that 50% of sentences for annotation include a specific point mutation. All together, this sampling provides better coverage of the different omic events and evidence types that were of interest. Special care is required when evaluating models built on this customized training set as an unweighted evaluation would not be representative of the real literature.

**Figure 2.**
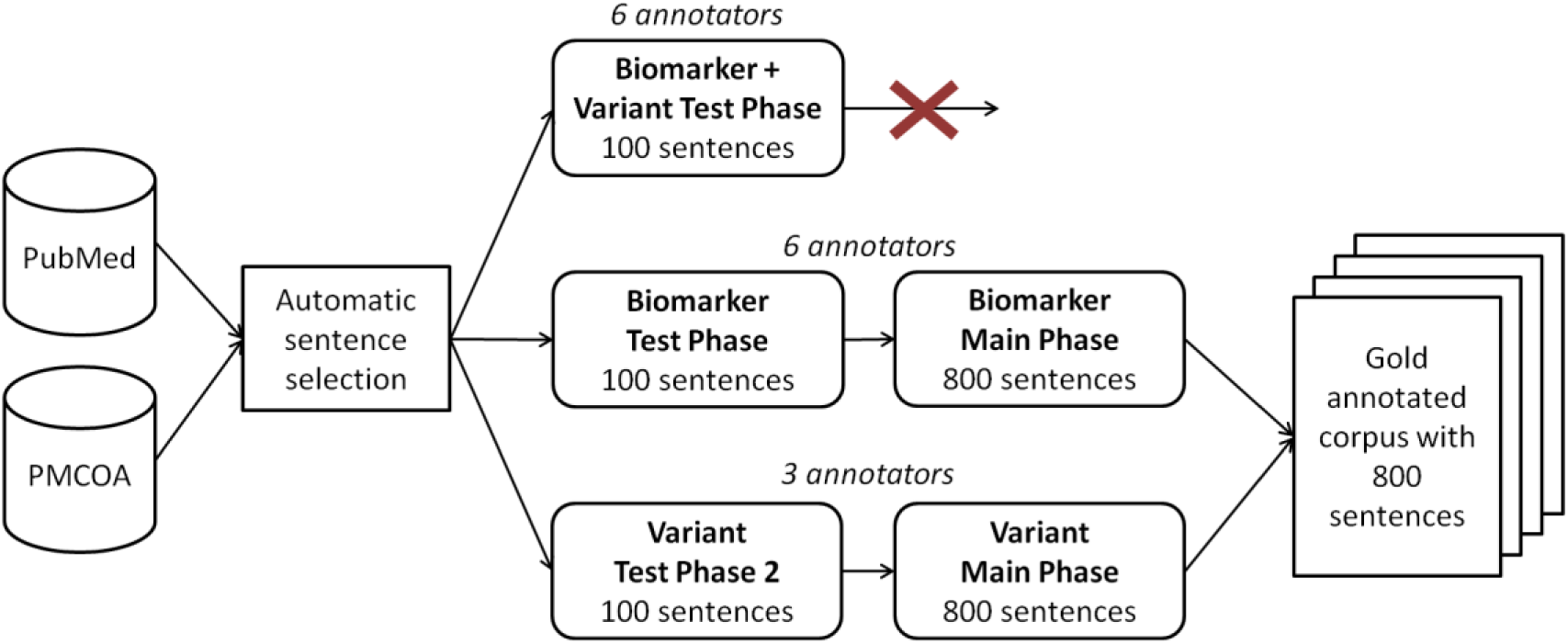
An overview of the annotation process. Sentences are identified from the literature that describe cancers, genes, variants and optionally drugs before being filtered using search terms. The first test phase tried complex annotation of biomarker and variants together but was unsuccessful. The annotation task was split into two separate tasks for biomarkers and variants separately. Each task had a test phase and then the main phase on the 800 sentences that were used to create the gold set.

Sentences that contain many permutations of relationships (e.g. a sentence with 6 genes and 4 cancer types mentioned) were removed. An upper limit of 5 possible relations was enforced for each sentence. This was done with the knowledge that the subsequent relation extraction step would have a greater false positive rate for sentences with very large number of possible relations. It was also done to make the annotation task more manageable. An annotation manual was constructed with examples of sentences that would and would not match the four evidence types. This was built in collaboration with CIViC curators and is available in our Github repository (https://github.com/jakelever/civicmine).

Each annotation task began with a test phase of 100 sentences. This allows the annotators to become accustomed to the annotation platform and make adjustments to the annotation manual to clarify misunderstandings. The first test phase (Biomarker + Variant) involved annotating sentences for ternary (gene, cancer, variant) or quaternary (gene, cancer, variant, drug) relationships. The ternary relationships included diagnostic, prognostic and predisposing and the quaternary relationship was predictive. A low F1-score inter-annotator agreement (average of 0.52) forced us to reconsider the annotation approach. From our evaluation, we determined that this poor agreement was likely due to including variants within the annotations. This created a large combinatorial problem where it was often unclear exactly which entity mentions to include within a relationship. In order to simplify the problem, the task was split into two separate annotation tasks, the biomarker annotation and the variant annotation. The biomarker annotation involved binary (gene, cancer) and ternary (gene, cancer, drug) relations that described one of the evidence types. The predictive and prognostic evidence types were merged (as shown in Figure 2), to further reduce the annotation complexity. The predictive/prognostic annotations could be separated after tagging as relationships containing a drug would be predictive and those without would be prognostic. A further postprocessing step to generate the gold set involved identifying prognostic relationships that overlapped with predictive relationships (i.e. shared the same gene and cancer type in a sentence) and removing them. The variant annotation task (gene, variant) focused on whether a variant (e.g. deletion) was associated with a specific gene in the sentence.

With the redefined annotation task, six annotators were involved in biomarker annotation, all with knowledge of the CIViC platform and having experience interpreting patient cancer variants in a clinical context. Three annotators (one of whom was involed in the biomarker annotation) were involved in variant annotationand they all had experience in cancer genomics. Both annotation tasks started with a new 100-sentence test phase to evaluate the redefined annotation tasks and resolve any ambiguity within the annotation manuals. Good inter-annotator agreement was achieved at this stage for both the biomarker annotation (average F1-score = 0.68) and variant annotation (average F1-score = 0.95). These 100 sentences were discarded as they exhibited a learning curve as annotators become comfortable with the task.

After a video-conference discussion, the annotation manuals were refined further. The main phase of biomarker annotation involved three annotators working on 400 sentences and the other three working on a different 400 sentences. Separately, three annotators worked on variant annotation with the 800 sentence set. Figure 3 shows the inter-annotator agreement for these tasks for the full 800 sentences. Each sentence is annotated by three annotators and a majority vote system is used to solve conflicting annotations. The biomarker and variant annotations are then merged to create the gold corpus of 800 sentences used for the machine learning system.

**Figure 3.**
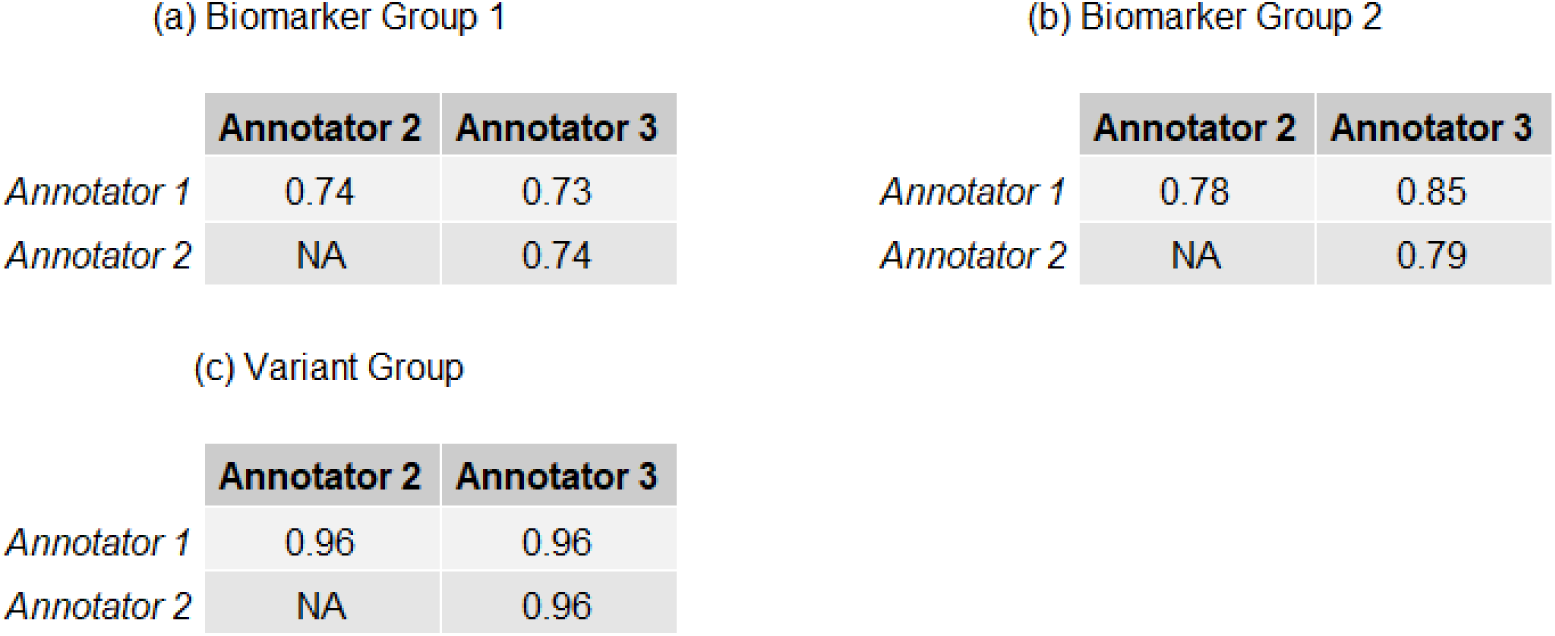
The inter-annotator agreement for the main phase for 800 sentences, measured with F1-score, showed good agreement in the two sets of annotations for biomarkers (a) and (b) as well as very high agreement in the variant annotation task (c). The sentences from the multiple test phases are not included in these numbers and were discarded from further analysis.

### Relation extraction

The sentences annotated with relations were then processed using the Kindred relation extraction Python package. Relation extraction models were built for all five of the relation types: the four evidence types (diagnostic, predictive, predisposing and prognostic) and one associated variant relation type. Three of the four evidence type relations are binary between a gene entity and a cancer entity. The associated variant relation type is also binary between a gene entity and a variant entity. The predictive evidence item type was ternary between a gene, a cancer type and a drug.

Most relation extraction systems focus on binary relations [34,35] and use features based on the dependency path between those two entities. The recent BioNLP Shared Task 2016 series included a subtask for non-binary relations (i.e. relations between three or more entities) but no entries were received [36]. Relations between 2 or more entities are known as n-ary relations where *n* ≥ 2. The Kindred relation extraction package, based on the VERSE relation extraction tool [37], which won part of the BioNLP Shared Task 2016, was enhanced to allow prediction of n-ary relations. First, the candidate relation builder was adapted to search for relations of a fixed *n* which may be larger than 2. This meant that sentences with 5 non-overlapping tagged entities would generate 60 candidate relations with *n* = 3. These candidate relations would then be pruned by entity types. Hence, for the predictive relation type (with *n* = 3), the first entity must be a cancer type, the second a drug and the third a gene. Two of the features used are based on the path through the dependency graph between the entities in the candidate relation. For relations with more than two entities, Kindred made use of a minimal spanning tree within the dependency graph. The default Kindred features were then constructed for this subgraph and the associated entities and sentences. All features were represented with 1-hot vectors or bag-of-words representations.

During training, candidate relations are generated with matching n-ary to the training set. Those candidate relations that match a training example are flagged as positive examples with all others as negative. These candidate relations are vectorized and a logistic regression classifier is trained against them. The logistic regression classifier outputs an interpretable score akin to a probability for each relation, which was later used for filtering. Kindred also supports a Support Vector Machine classifier (SVM) or can be extended with any classifier from the scikit-learn package [38]. The logistic regression classifier was more amenable to adjustment of the precision-recall tradeoff.

For generation of the knowledgebase, the four evidence type relations were predicted first which provided relations including a Gene. The Associated Variant relation was then predicted and attached to any existing evidence type relation that included that gene.

### Evaluation

With the understanding that the annotated sentences were selected randomly from customised subsets and not randomly from the full population, care was taken in the evaluation process.

First, the annotated set of 800 sentences was split 75%/25% into a training and test set that had similar proportions of the four evidence types (Table 2). Each sentence was then tracked with the group it was selected from (Table 1). Each group has an associated weight based on the proportion of the entire population of possible sentences that it represents. Hence, the prognostic group, which dominates the others, has the largest weight. When comparing predictions against the test set, the weighting associated with each group was then used to adjust the confusion matrix values. The goal of this weighting scheme was to provide performance metrics which would be representative for randomly selected sentences from the literature and not for the customised training set.

**Table 2.**
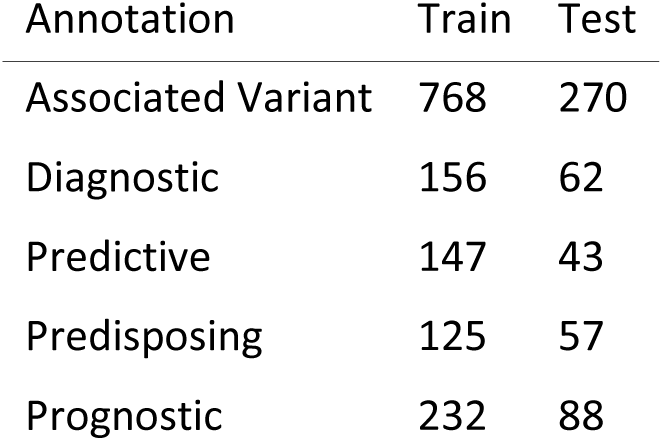
Number of annotations in the training and test sets

| Annotation | Train | Test |
| --- | --- | --- |
| Associated Variant | 768 | 270 |
| Diagnostic | 156 | 62 |
| Predictive | 147 | 43 |
| Predisposing | 125 | 57 |
| Prognostic | 232 | 88 |

### Precision-recall Tradeoff

Figure 4a shows precision recall curves for all five of the relation types. The diagnostic and predisposing tasks are obviously the most challenging for the classifier. This same data can be visualised by comparing the threshold values used against the output of the logistic regression for each metric (Fig 4b).

**Figure 4.**
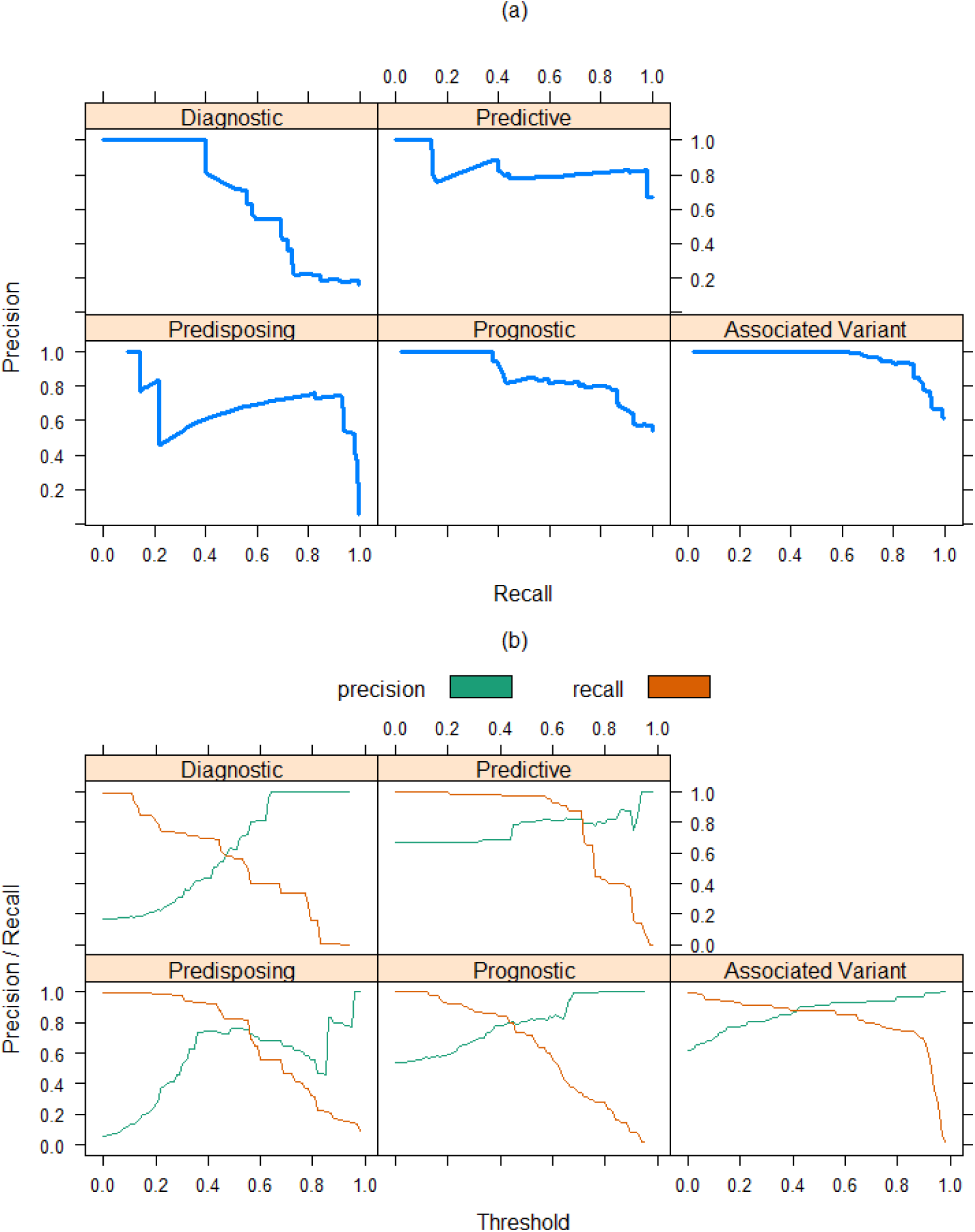
(a) The precision-recall curves illustrate the performance of the five relation extraction models built for the four evidence types and the AssociatedVariant prediction. (b) This same data can be visualised in terms of the threshold values on the logistic regression to select the appropriate value for high precision with reasonable recall.

In order to provide a high quality resource, we decided on a trade off of high precision with low recall. We hypothesised that the most commonly discussed cancer biomarkers, which are the overall goal of this project, would appear in many papers using different wording. These frequently mentioned biomarkers would then be likely picked up even with lower recall. This also reduces the burden on CIViC curators to sift through false positives. With this, we selected thresholds that would give as close to 0.9 precision given the precision-recall curves for the four evidence types. We require a higher precision for the variant annotation (0.94). The thresholds and associated precision recall tradeoffs are shown for all five extracted relations in Table 3.

**Table 3.**
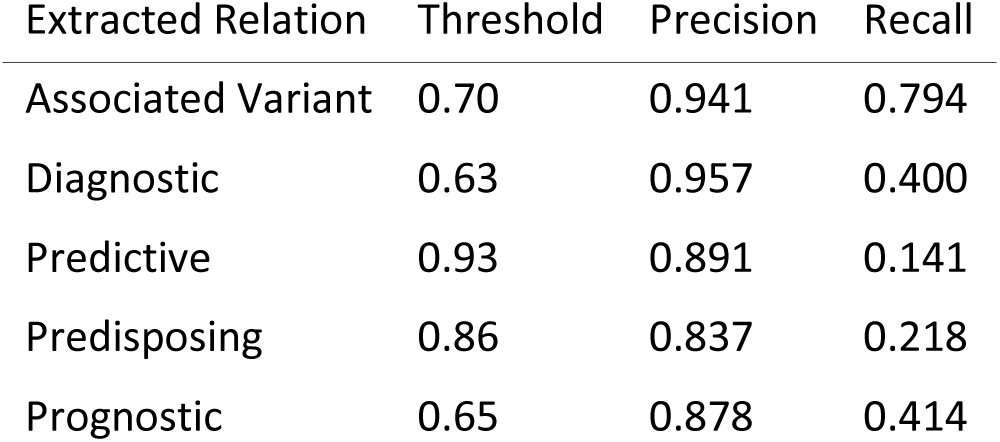
The selected thresholds for each relation type with the high precision and lower recall trade-off.

### Application to PubMed and PMCOA with Updates

With the thresholds selected, the final models were applied to all sentences extracted from PubMed and PMCOA. This is a reasonably large computational problem and was tasked to the compute cluster at the Genome Sciences Centre.

In order to manage this compute and provide infrastructure for easy updating with new publications in PubMed and PMCOA, we made use of the updated PubRunner infrastructure (paper in preparation - https://github.com/jakelever/pubrunner). This allows for easy distribution of the work across a compute cluster. The resulting data was then pushed to Zenodo for perpetual and public hosting (https://doi.org/10.5281/zenodo.1156241). The data is released with a Creative Commons Public Domain (CC0) license so that other groups can easily make use of it.

The PubRunner infrastructure enables the easy update of the resource. We plan to update the resource every month. It manages download and execution of the tool as well as the upload of the data to the Zenodo repository.

### CIViC Matching

In order to make comparisons with CIViC, we downloaded the nightly data file from CIViC (https://civicdb.org/releases – downloaded on 03 December 2018) and matched evidence items against items in CIViCmine. The evidence type and IDs for genes and cancers were used for matching. Direct string matching was used to compare drug names for predictive biomarkers. The exact variant was not used for comparison in order to find genes that contain any biomarkers that match between the two resources.

Some mismatches occurred with drug names. For example, CIViCmine may capture information about the drug family while CIViC contains information on specific drugs, or a list of drugs. Another challenge with matching with CIViCmine is related to the similarity of cancer types in the Disease Ontology. There are several pairs of similar cancers types that are used interchangably by some researchers and not by others, e.g. stomach cancer and stomach carcinoma. CIViC may contain a biomarker for stomach cancer and CIViCmine matches all the other details except it relates it to stomach carcinoma.

### User interface

In order to make the data easily explorable, we provide a Shiny-based front-end (Fig 5) [39]. This shows a list of biomarkers extracted from abstracts and papers, which can be filtered by the Evidence Type, Gene, Cancer Type, Drug and Variant. In order to help prioritize the biomarkers, we use the number of unique papers that the variants are mentioned in as a metric. By default, the listed biomarkers are shown with the highest citation count first. Whether the biomarker is found in CIViC is also shown as a column and is an additional filter. The CIViC information is updated daily by downloading the latest nightly release. This allows CIViC curators to quickly navigate to biomarkers not currently discussed in CIViC and triage them efficiently.

**Figure 5.**
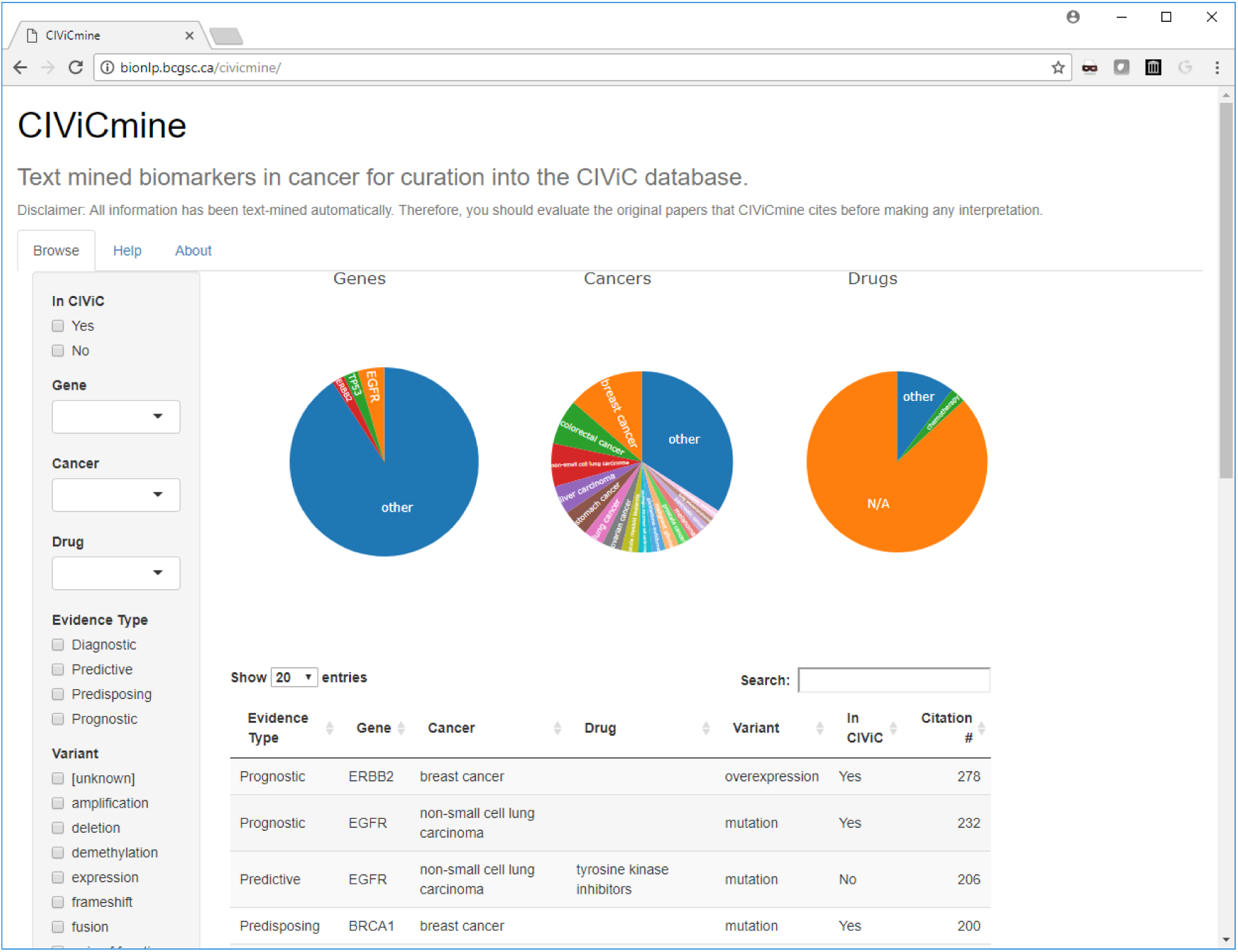
A Shiny-based web interface allows for easy exploration of the CIViCmine biomarkers with filters and overview piecharts. A main table shows the list of biomarkers and links to a subsequent table showing the list of supporting sentences.

With filters selected, the user is presented with pie-charts that illustrate the representation of different cancer types, genes and drugs. When the user clicks on a particular biomarker, an additional table is populated with the citation information. This includes the journal, publication year, section of the publication (e.g. title, abstract or main body), subsection (if cited from the main body) and the actual text of the sentence from which the relationship was extracted. This table can further be searched and sorted, for example, to look for older citations or citations from a particular journal. The PubMed ID is also provided with a link to the citation on PubMed.

## Results

From the full PubMed and PMCOA corpus, we extracted 90,992 biomarkers with a breakdown into the four types (Figure 6). As expected, based on our preliminary analysis, there are many more prognostic evidence items than the other three types. Table 4 outlines examples of all four of these evidence types. 36.4% of sentences (46,931/128,857) contain more than one evidence item, such as the predictive example which relates *EGFR* as a predictive marker in NSCLC to both erlotinib and gefitinib. In total, we extracted 202,390 mentions of biomarkers from 69,258 unique papers. These biomarkers relate to 7,866 genes, 557 cancer types and 402 drugs.

**Table 4.**
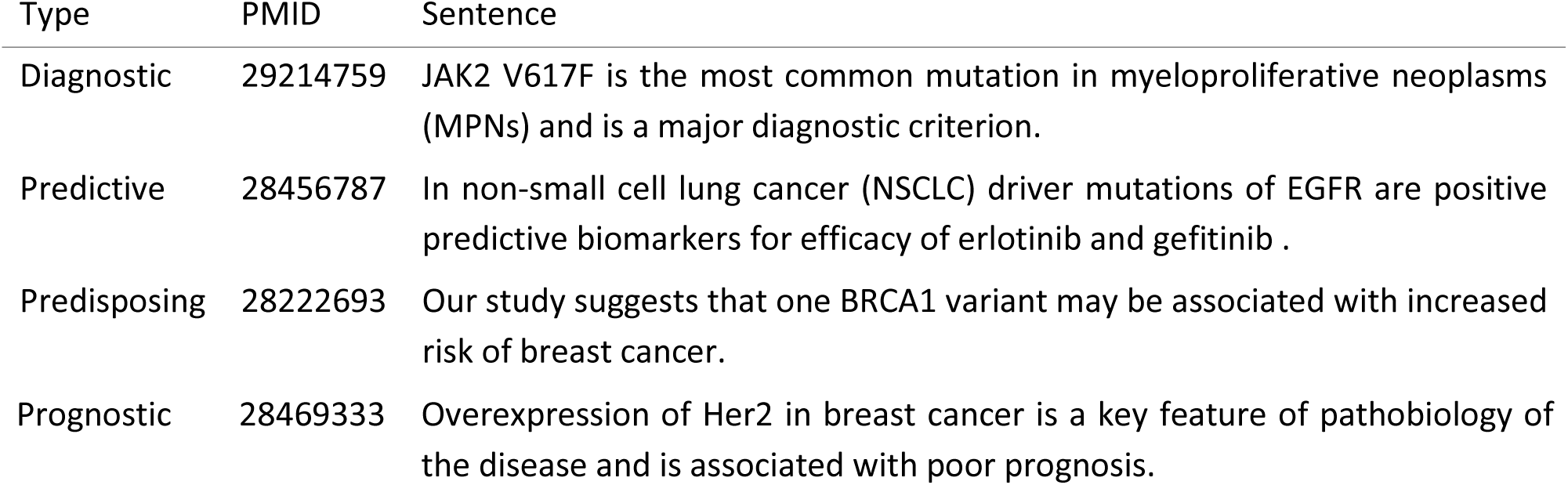
Four example sentences for the four evidence types extracted by CIViCmine. The associated PubMed IDs are also shown for reference.

**Figure 6.**
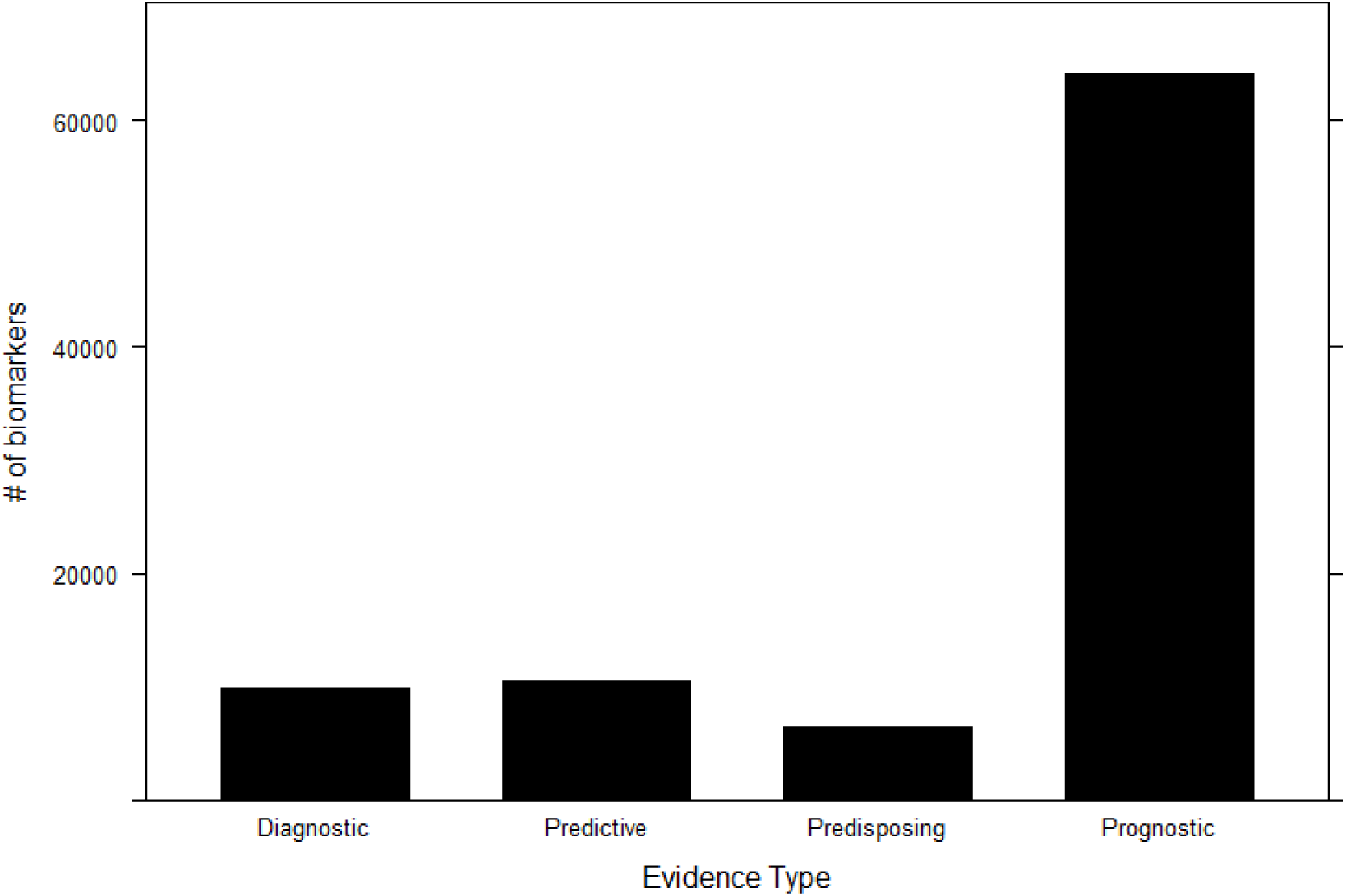
The entirety of PubMed and PubMed Central Open Access subset were processed to extract 90,992 biomarkers distributed between the four different evidence types shown.

*EGFR* and *TP53* stand out as the most frequently extracted genes in different evidence items (Fig 7a). Over 50% of the *EGFR* evidence items are associated with lung cancer or non-small cell lung carcinoma (NSCLC). *CDKN2A* has a larger proportion of diagnostic biomarkers associated with it than most of the other genes in the top 20. *CDKN2A* expression is a well-established marker for distinguishing HPV+ versus HPV-cervical cancers. Its expression or methylation states are discussed as diagnostic biomarkers in a variety of other cancer types including colorectal cancer and stomach cancer.

**Figure 7.**
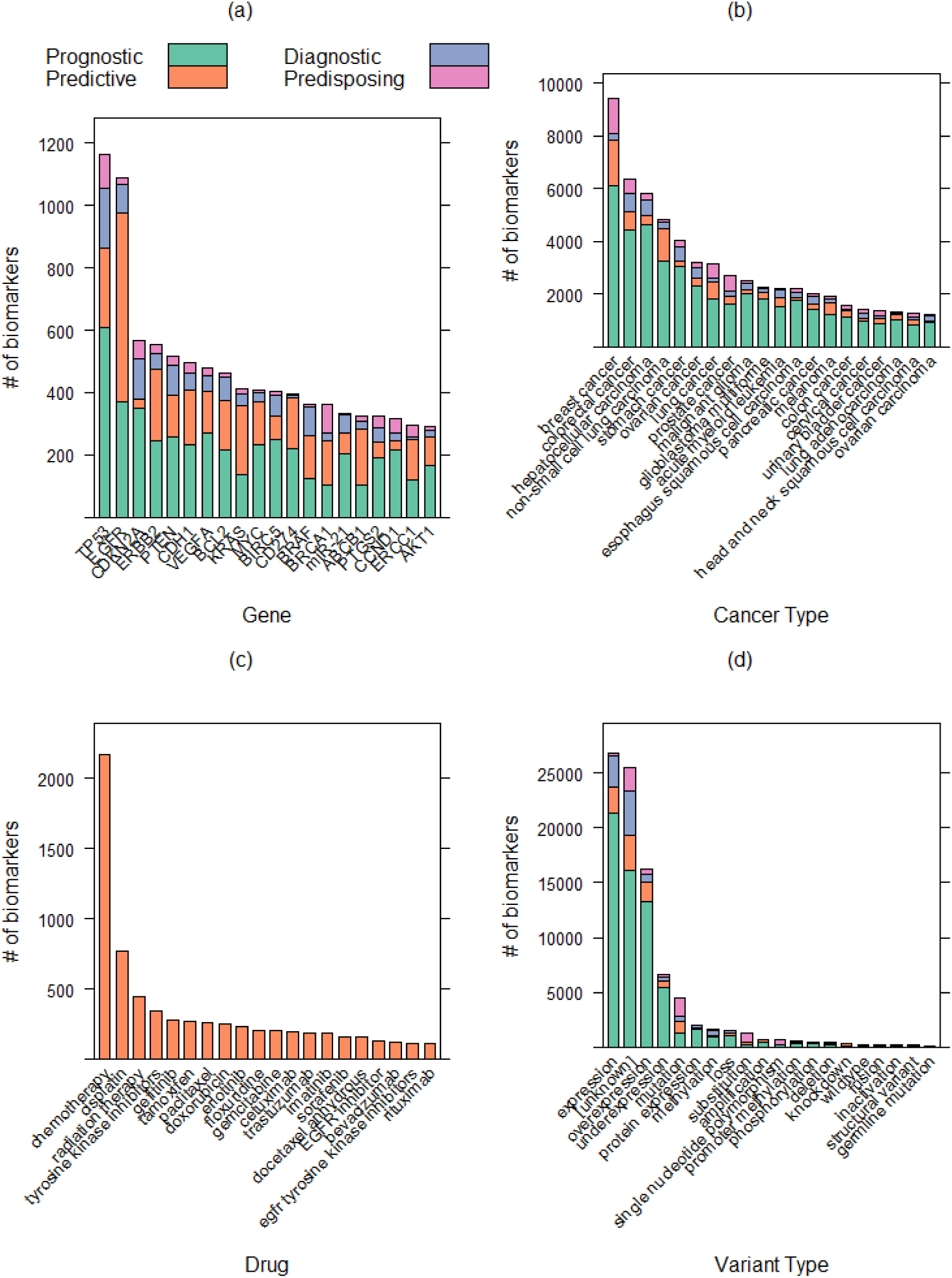
An overview of the top 20 (a) genes, (b) cancer types, (c) drugs and (d) variants extracted as part of evidence items.

Breast cancer is, by far, the most frequently discussed cancer type (Fig 7b). A number of the associated biomarkers focus on predisposition, as breast cancer has one of the strongest hereditary components associated with germline mutations in *BRCA1* and *BRCA2*. NSCLC shows the largest relative number of predictive biomarkers, consistent with the previous figure showing the importance of *EGFR*.

For the predictive evidence type, we see a disproportionally large number associated with the general term chemotherapy and specific types of chemotherapy including cisplatin, paclitaxel and doxorubicin (Fig 7c). Many targeted therapies are also frequently discussed such as the *EGFR* inhibitors, gefitinib, erlotinib and cetuximab. More general terms such as “tyrosine kinase inhibitor” capture biomarkers related to drug families.

Lastly, we see that expression related biomarkers dominate the variant types (Fig 7d). Markers based on expression are more likely to be prognostic than those using non-expression data (80.7% versus 44.4%). The popular approach to exploring the importance of a gene in a cancer type is to correlate expression levels with patient survival. With the extended historical use of immunohistochemical methods as well as the accessibility of large transcriptome sets and survival data (e.g. TCGA), such associations have become very common. The ‘mutation’ variant type has a more even split across the four evidence types. The mutation term covers very general phrasing without a specific mention of a specific mutation. The substitution variant type does capture this information but there are far fewer than biomarkers with the ‘mutation’ variant type. This reflects the challenge of extracting all of the evidence item information from a single sentence. It is more likely for an author to define a mutation in another section of the paper or aggregate patients with different mutations within the same gene and then use a general term (e.g. *EGFR* mutation) when discussing its clinical relevance. There are also a substantial number of evidence items where the variant cannot be identified and are flagged as ‘[unknown]’. These are still valuable but may require more in-depth curation in order to identify the actual variant.

Of all the biomarkers extracted, 21.9% (19,957/ 90,992) are supported by more than one citation. In fact, the most cited biomarker is *BRCA1* mutation as a predisposing marker in breast cancer with 725 different papers discussing this. The initial priority for CIViC annotation is on highly cited biomarkers that have not yet been curated into CIViC, in order to eliminate obvious information gaps. However, the single citations may also represent valuable information for precision cancer analysts and CIViC curators focused on specific genes or diseases.

We compared the 90,992 biomarkers extracted by CIViCmine with the 2,380 in the CIViC resource as of 03 December 2018. Figure 8a shows the overlap of exact evidence items between the two resources. The overlap is quite small and the number evidence extracted in CIViCmine not yet included in CIViC is very large. We next compare the cited publications using PubMed ID. Despite not having used CIViC publications in training CIViCmine, we find that a substantial number of papers cited in CIViC (316/1,425) were identified automatically by CIViCmine (Fig 8b). The remaining ∼1100 papers were likely not identified as they did not contain a single sentence that contained all the information necessary for extraction. Future methods that can identify biomarkers discussed across multiple sentences would likely identify more of these papers. Altogether, CIViCmine includes 6,449 genes, 432 cancer types and 333 drugs or drug families not yet included in CIViC.

**Figure 8.**
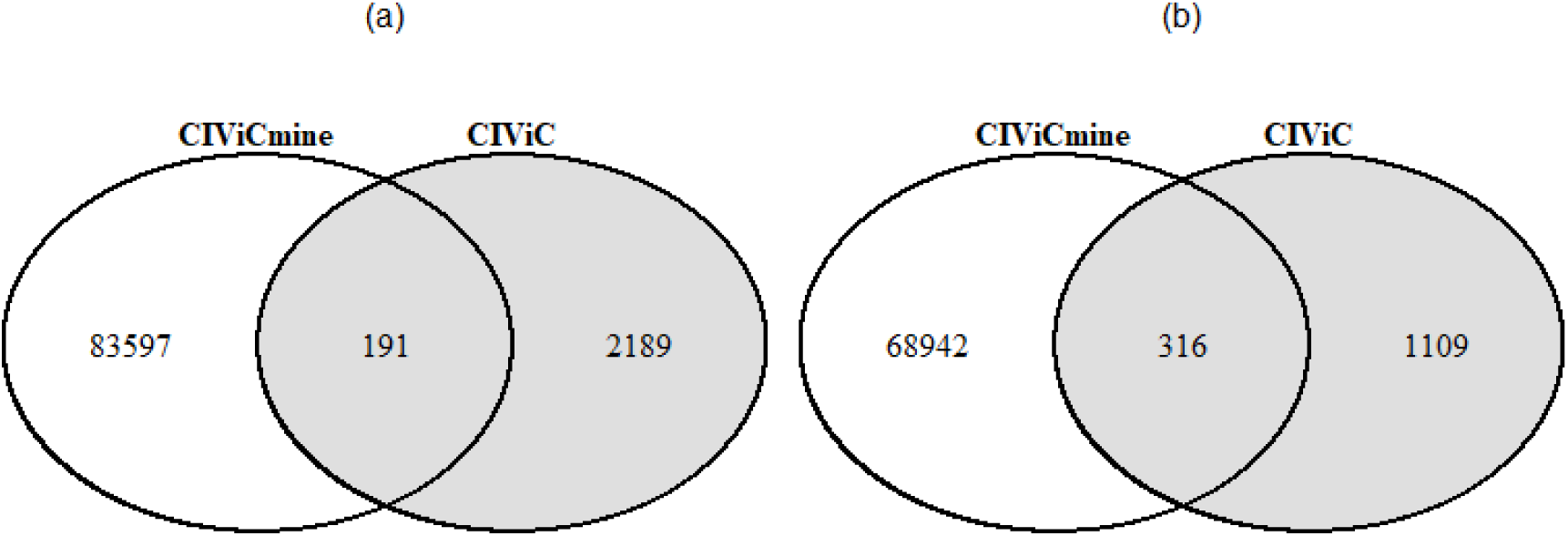
A comparison of the evidence items curated in CIViC and automatically extracted by CIViCmine by (a) exact biomarker information and by (b) paper.

### Use Cases

There are two use cases of this resource that are already been realised by CIViC curators at the McDonnell Genome Institute and analysts at BC Cancer.

Knowledge-base curation use case: The main purpose of this tool is to assist in curation of new biomarkers in CIViC. A CIViC curator, looking for a frequently discussed biomarker, would access the CIViCmine Shiny app through a web browser. This would present the table, pie charts and filter options on the left. They would initially filter the CIViCmine results for those not already in CIViC. If they had a particular focus, they may filter by Evidence Type. For example, some CIViC curators may be more interested in diagnostic, predictive and prognostic biomarkers than predisposing. This is due to the relative importance of somatic events in many cancer types. They would then look at the table of biomarkers, already sorted by citation count in descending order, and select one of the top ones. This would then populate a table further down the page. Assuming that this is a frequently cited biomarker, there would be many sentences discussing it, which would quickly give the curator a broad view of whether it is a well-supported association in the community. They might then open multiple tabs on their web browser to start looking at several of the papers discussing it. They might select an older paper, close to when it was first established as a biomarker, and a more recent paper from a high-impact journal to gauge the current view of the biomarker. Several of the sentences may obviously cite other papers as being important to establishing this biomarker. The curator would look at these papers in particular, as they may be the most appropriate to curate. Importantly, the curator can use this to identify the primary literature source(s), which includes the experimental data supporting this biomarker.

Personalized cancer analyst use case: While interpreting an individual patient tumor sample, an analyst typically needs to interpret a long list of somatic events. Instead of searching PubMed for each somatic event, they can initially check CIViC and CIViCmine for existing structured knowledge on the clinical relevance of each somatic event. First, they should check CIViC given the high level of pre-existing curation there. This would involve searching the CIViC database through their website or API. If the variant does not appear there, they would then progress to CIViCmine. By using the filters and search functionality, they could quickly narrow down the biomarkers for their gene and cancer type of interest. If a match is found, they can then move to the relevant papers that are listed below to understand the experiments that were done to make this assertion. As they evaluate this biomarker, they could enter this evidence and all of the structured fields that may be spread throughout the publication into the CIViC database. Both CIViC and CIViCmine reduce curation burden by aggregating likely applicable data across multiple synonyms for the gene, disease, variant or drug not as easily identified through PubMed searches.

### Evaluation by CIViC curator

In order to evaluate the curation value of the data provided by CIViCmine, a CIViC curator evaluated the top biomarkers identified by CIViCmine that were not found in CIViC. Biomarkers with high citation counts were selected for each evidence type and filtered for those which the variant was also extracted. They were then evaluated for correctness (whether the sentences matched the extracted structured data), usability (whether there was enough information for curation into CIViC contained within the sentence) and need (whether this information was lacking in CIViC). Each biomarkers was marked in all three categories with Yes, Intermediate and No. Intermediate scores are used to identify cases where additional information (e.g. reading the full paper or its citations) was needed. Figure 9 shows the summary of the results as percentages for each of the three metrics across the four evidence types.

**Figure 9.**
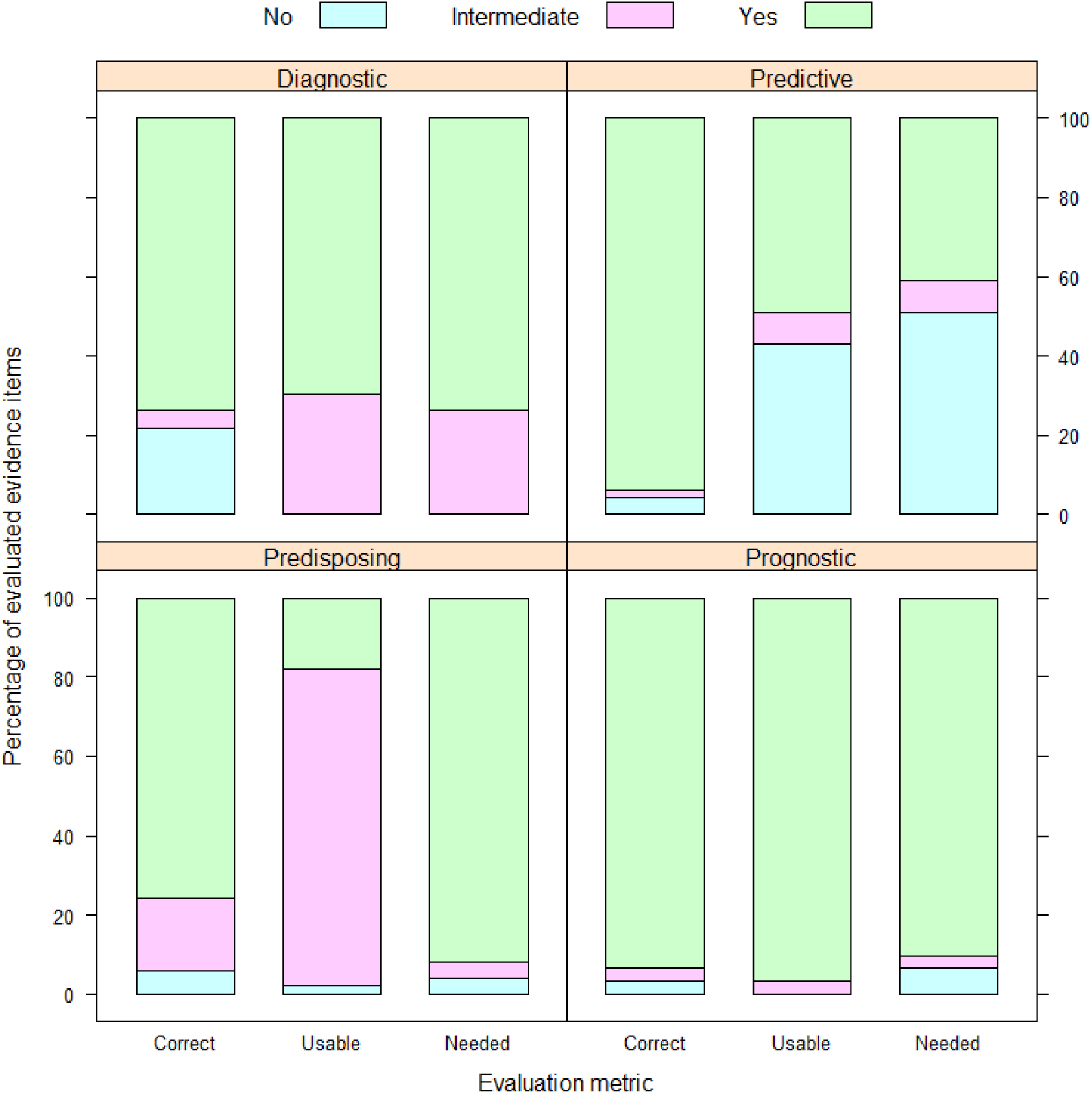
The top results in CIViCmine were evaluated by a CIViC curator and measured for three categories (correctness, usability and need). Percentages are shown for each metric and evidence type for No, Intermediate and Yes.

Overall, the results are very positive with 73% of evaluated biomarkers being deemed needed by CIViC. The predictive evidence type was found to have a larger proportion of unneeded evidence items. This was due to the catch all groups (e.g. *EGFR* inhibitors) that were deemed to be too vague for inclusion into CIViC but might provide valuable information for other clinical researchers. The high percentage of Intermediate for the usability of predisposing biomarkers was due to the general variant terms identified (such as Mutation) where the exact variant was unclear and further curation would be needed. Overall, these results show that CIViCmine provides valuable data that can be curated into CIViC and other knowledgebases.

## Discussion

This work provides several significant contributions to the fields of biomedical text mining and precision oncology. Firstly, the annotation method is drastically different from previous approaches. Most annotation projects (such as the BioNLP Shared Tasks [40,41] and the CRAFT corpus [42]) have focused on abstracts or entire documents. The biomarkers of interest for this project appear sparsely in papers so it would have been inappropriate to annotate full documents and a focus on individual sentences was necessary. In selecting sentences, we aimed for roughly half the sentences to contain positive relations. This would enable better classifier training with a more even class balance. Therefore, we filtered the sentences with a series of keywords after identifying those that contain the appropriate entities This approach could be applied to many other biomedical topics.

We also made use of a simpler annotation system than the often used brat [43] which allowed for fast annotation by restricting the possible annotation options. Specifically, annotators did not select the entities but were shown all appropriate permutations that matched the possible relation types. Issues of incorrect entity annotation were reported through the interface, collated and used to make improvements to the underlying wordlists for gene, cancer types and drugs. We found that once a curator became familiar with the task, they could curate sentences relatively quickly with approximately 1-2 minutes spent on each sentence. Expert annotation is key to providing high quality data to build and evaluate a system. Therefore reducing the time required for expert annotators is essential.

The supervised learning approach differs from methods that used co-occurrence based (e.g. STRING [25]) or rule-based (e.g. mirTex [24]) methods. Firstly, the method is able to extract complex meaning from the sentence providing results that would be impossible with a co-occurrence method. A rule-based method would require enumerating the possible ways of describing each of the diverse evidence types. Our approach is able to capture a wide variety of biomarker descriptions. Furthermore, most relation extraction methods aim for optimal F1-score [36], placing an equal emphasis on precision and recall. With the goal of minimizing false positives, our approach of high precision and low recall would be an appropriate model for other information extraction methods applied to the vast PubMed corpus.

Apart from the advantages outlined previously, several other factors lead to the decision to use a supervised learning approach to build this knowledgebase. The CIViC knowledgebase could have been used as training data in some form. The papers already in CIViC could have been searched for the sentences discussing the relevant biomarker, which could then have been used to train a supervised relation extraction system. An alternative approach to this problem would have been to use a distant supervision method using the CIViC knowledgebase as seed data. This approach was taken by *Peng et al*, who also attempted to extract relations across sentence boundaries [44]. They chose to focus only on point mutations and extracted 530 within sentence biomarkers and 1,461 cross-sentence biomarkers. These numbers are drastically smaller than the 70,655 extracted in CIViCmine.

The reason to not use the CIViC knowledgebase in the creation of the training data was taken to avoid any curator-specific bias that may have formed in the selection of papers and biomarkers already curated. Such biases within the CIViC knowledgebase are already known, common within various clinical knowledgebases, and driven by collaborations with experts in specific fields or institutional expertise. Avoiding this bias was key to providing a broad and unbiased view of the biomarkers discussed in the literature. CIViC evidence items include additional information such as directionality of a relationship (e.g. does a mutation cause drug sensitivity or resistance), the level of support for it (from preclinical models up to FDA guidelines) and several other factors. It is highly unlikely that all this information will be included within a single sentence. Therefore, we did not try to extract this information concurrently. Instead, it is an additional task for the curator as they process the CIViCmine prioritised list.

A robust named entity recognition solution does not exist for a custom term list of cancer types, drugs and variants. For instance, the DNorm tool does not capture many cancer subtypes. A decision was made to go for high recall for entity recognition, including genes, as the relation extraction step would then filter out many incorrect matches based on context. This decision is further supported by the constant evolution of cancer type ontologies, as demonstrated by workshops at recent Biocuration conferences.

The CIViCmine app, accessible at http://bionlp.bcgsc.ca/civicmine, and freely accessible associated data provide a valuable addition to the precision oncology informatics community. CIViCmine can be used to assist curation of other precision cancer knowledgebases and can be used directly by precision cancer analysts to search for biomarkers of interest. As this resource will be updated monthly with the latest research, it will constantly change as new cancer types and drug names enter the lexicon. We anticipate that the methods described can be used in other biomedical domains and that the resources provided will be valuable to the biomedical text mining and precision oncology fields.

## Acknowledgements

CIViC is supported by the National Cancer Institute (NCI) of the National Institutes of Health (NIH) under award number U01CA209936. JL is supported by a Vanier Canada Graduate Scholarship. MG is supported by the NHGRI under award number R00HG007940.

